# Third-party imitation is not restricted to humans

**DOI:** 10.1101/2025.03.14.643244

**Authors:** Esha Haldar, Ariana Hernandez Sanchez, Claudio Tennie, Sara Torres Ortiz, Janneke Vos, Maurice Valbert, Auguste M.P. von Bayern

## Abstract

Imitation of cultural practices is ubiquitous in humans and often involves faithful copying of intransitive (i.e., non-object directed) gestures and societal norms which play a crucial role in human cultural evolution. Apart from learning these directly from a tutor, humans often learn passively as third-party observers from the interactions of two or more individuals. Whether third-party imitation has evolved outside humans remains unknown. In the current study, we investigated whether undomesticated blue-throated macaws could imitate in a third-party setting. A naïve test group (N=6) passively observed a conspecific demonstrator performing improbable intransitive actions in response to specific human gestural commands. Directly afterwards, the observer received the same gestural commands and performance-contingent rewards. An equally naïve control group (N=5) was tested correspondingly, in the absence of third-party demonstrations. The test group learned more target actions (mean=4.16 versus mean=2.2) in response to the specific commands, significantly faster and performed them more accurately than the control group. The test group also spontaneously imitated some of the actions even before they received any gestural commands or rewards. Our findings show that third-party imitation, even for intransitive actions, exists outside humans, allowing for rapid adaption to group specific behaviours and possibly cultural conventions in parrots.

## INTRODUCTION

Human cultural evolution is largely underpinned by the human ability to imitate. While a large part of human culture comprises the learning of technical skills involving tools and artefacts, another substantial part requires the learning of arbitrary cultural conventions ^1,2^ encompassing intransitive actions (i.e., actions not directed at objects) like gestures ^3^ or dance movements ^4^. Humans can quickly adopt group-specific behavioural norms (which may eventually lead to cultural conventions), by learning directly from overt demonstrations of a skilled individual ^5^ or by passively observing skilled individuals without any direct interaction ^6,7^. Research has shown that in many small-scale cultures, particularly extant hunter-gatherer societies, learning by passive observation is more prevalent than learning via overt demonstration (teaching) ^6–8^. Children in such societies learn technical skills like tool use, hunting etc. by passively observing skilled adults ^9^ or learn about social interaction behaviours, language etc. by attending to interaction of two or more group members ^10,11^ as third-party observers. While human children develop the tendency to imitate an adult who is directly demonstrating an action to the child (A learns from B), from the first year of their life, they develop such third-party imitation (A learns from B and C interacting) ability only towards the end of their second year ^12–15^, once they have developed an understanding for self-other equivalence ^13,16^. Thus, third-party imitation is an important skill that has been reported only in humans ^17^ and the evolution of this ability remains largely unexplored. A comparative approach investigating third-party imitation in non-human, non-primate species promises to shed light on whether this ability evolved convergently in other distant animal taxa.

Third-party imitation entails learning of an act passively, by observing the interaction of two individuals ^13^. Passive observational learning by imitation of a single demonstrator has been reported in many species ^18–21^, but no study has reported imitation in a third-party context. In order to investigate a non-human animal species’ ability to learn via third-party imitation in an ecologically valid manner, it is vital to ensure that test subjects (observers) are neither human-trained in target actions nor in imitation/action copying (such as in ‘do-as-I-do’ or DAID studies; ^22–27^ nor must they be human-enculturated (like dogs ^28^). Also, their observation of the demonstrations should not be human directed (e.g., by using ostensive signals like calling, or eye gaze). Most studies investigating imitation in animals have either not fulfilled these criteria or did not investigate third-party imitation specifically.

As said initially, a substantial part of human culture (besides imitating transitive actions, e.g., when learning to manipulate objects or use tools) comprises the learning of intransitive actions, e.g., gestures, which often occurs via third-party imitation. Although gestural communication is found to be ubiquitous in non-human primate species ^29,30^, gestures are typically either innate responses ^31^ or ritualized ^32,33^, but do not seem to be acquired through imitation (neither through direct nor third party imitation). Moore ^34^ reported direct imitation of intransitive gestures in an African Grey parrot (*Psittacus erithacus*). The experimenters interacted directly with the parrot several times a day, performing certain stereotyped body movements. Over the course of five years, the parrot produced 10 actions using head, beak, wings and feet and was claimed to have mimicked intransitive actions of the human demonstrators. The other few studies that have reported imitation of intransitive actions in animals have used the “Do-as-I-do” (DAID) or “Do-as-the-other-does” (DATOD) paradigm, in which the animals are actively trained by the experimenter to pay attention to and copy either a human or conspecific demonstrator (^22–26^. All these studies were conducted in a two-party setting, and thus did not qualify for third party imitation.

Studies examining third-party imitation and its development in human children have assessed the learning performance of children when passively observing the interaction between a demonstrator and another individual (child A learning from B and C). Children aged 15-20 months old successfully learned about tools and object manipulation via third-party imitation ^12–14^. However, in non-human animals, only one single study has tested third party imitation in untrained individuals, namely a study on dogs (*Canis familiaris)*, a domesticated species ^17^. Here, a trained dog performed two actions in response to a human trainer giving verbal commands, while a task-naïve, non-DAID trained observer dog (i.e., the “third-party”) watched the interaction. Subsequently, it was tested whether the observer dog would itself show the actions demonstrated by the trained conspecific. None of the N=202 dogs tested produced the target actions with this procedure. Thus, dogs failed to evidence spontaneous third-party imitation in this study. Yet, even if some dogs had been successful, the obtained results could have been confounded by domestication effects in dogs ^17^. In a different set of studies, Pepperberg and colleagues, used a method called “model/rival technique” ^35^ in which two trainers interacted with each other while a single African grey parrot observed their interaction ^36^. However, this technique was employed as a general tool for training the parrot to learn object labels by vocal imitation, rather than for specifically investigating its ability to learn via third-party imitation. Moreover, the grey parrot received direct interactive training concurrently and hence the effect of learning as a third party cannot be teased apart retrospectively from these experiments. Thus, third-party imitation in non-human animals has not been studied specifically, even though a comparative approach promises insights into the evolution of third-party imitation. Given the previous studies^34,36^ indicating that parrots may be interesting candidates for studying third party imitation, we opted to investigate an undomesticated parrot species-blue-throated macaws (*Ara glaucogularis*).

Parrots are among the very few animal taxa possessing vocal imitation ability ^37–39^, tool-making skills ^40,41^, foraging cultures ^42,43^ and vocal dialects ^44^. They have relatively large brains with a high density of neurons ^37^ that seem to correspond to their advanced behavioural and cognitive complexity. Perhaps most importantly, parrots are impressive social learners ^40^ and have previously been successful in direct imitation of transitive and intransitive actions ^18,34,45^ but it remains untested whether they are capable of learning via third-party imitation (especially of intransitive actions).

We operationally defined third-party imitation as an increase in the likelihood to express otherwise improbable target actions upon observing them in a third-party setting, i.e., observing the interaction of a conspecific demonstrator and an experimenter. This improbability was established through a baseline data collection on the natural occurrence/frequency of our target actions and post-hoc analysis. In the experiment, a test group of n=6 hand-raised blue-throated macaws that were habituated to receiving food from human hands, but were not trained to perform specific behaviours on command and were never intentionally trained to imitate, observed a trained conspecific demonstrator performing five intransitive target actions (lift leg, fluff, spin, vocalise and flap wings), one at a time, in a session. Each target action was performed in response to specific hand commands from the experimenter (see Supplementary Information Figure S1). Subjects in the test group observed these demonstrations passively from an adjacent room separated by a transparent plexiglass (see Figure 1). Following these demonstrations (i.e., after 3-5 seconds), a second experimenter (who had been passively present in the room together with the subject) gave the same gestural commands to the subject (see Supplementary Videos S1-5), testing whether subjects would express the demonstrated target actions in 12 (±2) seconds of response time. A control group (N=5) was exposed to the same gestural commands, yet without ever observing any target action linked to the commands (see Supplementary Video S6-7). We rewarded the subjects in both test and control group for performing the target action within 12 (±2) sec following the respective command. If a subject performed the matching action simultaneously with the demonstration or before it received its own gestural command, it was not rewarded.

**Figure 1.**
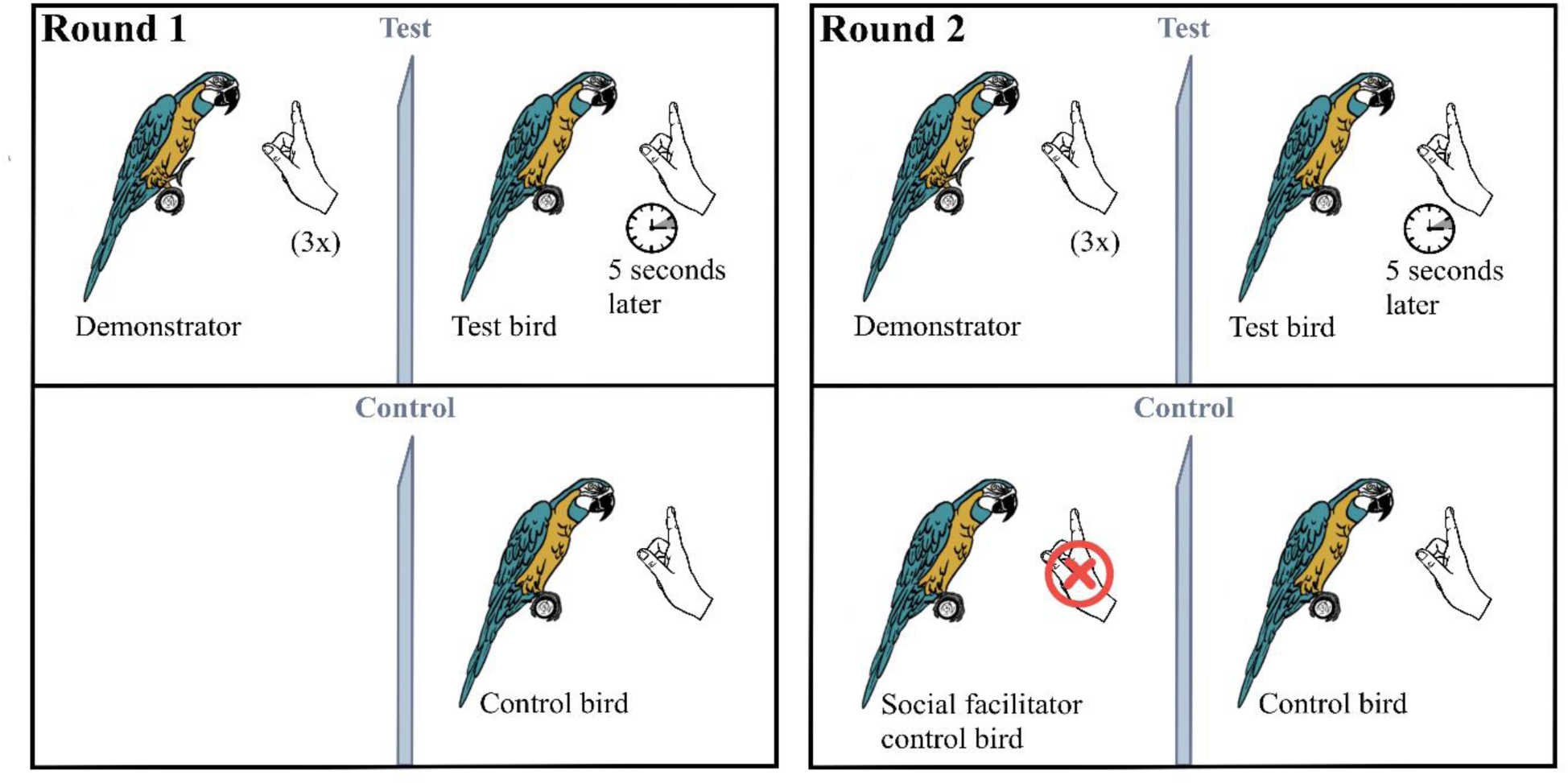
Illustration of the experimental procedure. The left panel shows the first round of testing, with the demonstrator and the test subject in the test group (left up) and the control subject tested alone in the control group (left down). The demonstrator performs an action thrice in response to the gestural command of the experimenter, 3-5 seconds following which the second experimenter gives the same gestural command to the test subject in a test trial. The control subject is only given the gestural command without any demonstration. In round 2, shown on the right panel, the testing contingency remains the same except for the addition of a social facilitator bird in the adjacent room of the control subject, who sits on the perch passively without performing any action.

## RESULTS

### Baseline: Establishing natural occurrence rates of target actions

A baseline of the natural occurrence of the chosen intransitive actions *‘lift leg’ ‘fluff’, ‘spin’, ‘vocal’* and *‘flap wings’* (see Supplementary Information Table S1) was established by conducting 4 hours of focal behavioural sampling of 11 subjects (44 hours in total) (1 control was unavailable) in the aviaries before the onset of the experiment. Behavioural observations throughout 15 min per subject were conducted for 16 days (resulting in a total of 4h in total per individual). From the yielded cumulative scores for each action (see Supplementary Information), we obtained the rate at which they occur in 12 sec response interval we gave to our subjects in the test and control conditions (see above). The actions were considered very improbable to occur naturally (according to criteria set post-hoc) within the response time except *‘vocal’* occurring infrequently (5%). The rate at which the other actions occurred naturally: *‘fluff’*: 0.5%; *‘flap wings’*: 0.04%; *‘spin’*: 0.001% and *‘lift leg’* 0%. *Lift leg* never occurred unless the bird scratched, nibbled its toes or fended of an aggressive approach by holding out a leg (which happened 0.03%) (see SM for details).

### Number of actions learnt

The subjects in the test condition expressed significantly more target actions upon commands than subjects in the control condition (Wilcoxon-rank-sum-test: W = 25, p = .005; effect size of r = .933, 95% CI [.794/∞]). A single, 7th subject from the test group did not learn any action. She had only recently been integrated into the group of our test birds at the age of 15 and did not socialize with the other individuals in the aviary for the entire run of the experiment, behaving passively throughout. We concluded that she had not yet sufficiently accommodated well to the group, the lab and testing contingencies. On these grounds she was removed from the analysis, reducing the sample size to 6 in the test condition. The 6 birds in the test group learnt on average 4 out of 5 actions (Mean=4.16, SE=±0.4) whereas the control group learnt 2 (Mean=2.20, SE=±0.2) as illustrated in Figure 2. Table 1 gives an overview of the group performance while individual performance for both groups is provided in Supplementary Information Table S2.

**Figure 2.**
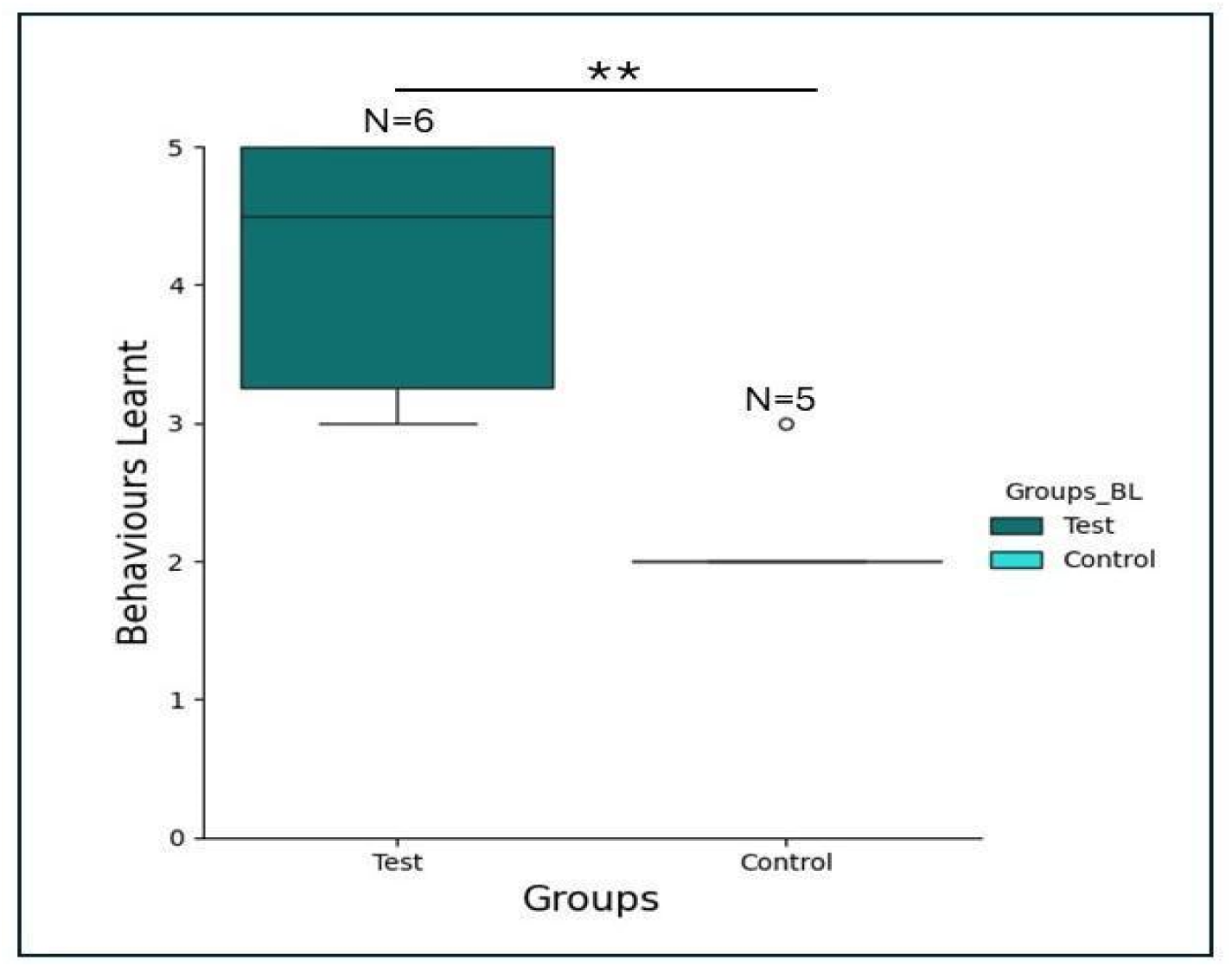
Number of target actions learnt by the subjects in the two experimental groups. The box plots show that the subjects in the test condition (teal blue) performed significantly better than the control group, denoted by asterisks (Wilcoxon-rank-sum-test (W = 25, p = .005). Error bars, mean ± SEM.

**Table 1.** Number of subjects in the test and control conditions who learnt the behaviours.

| Actions | Test (N=6 excluding a subject) | Control (N=5) |
| --- | --- | --- |
| Lift leg | N=6 | N=3 |
| Spin | N=6 | N=4 |
| Fluff | N=5 | N=3 |
| Vocal | N=5 | N=1 |
| Wings | N=3 | N=0 |

### Response accuracy

Mean correct response rate for each subject was calculated as the total number of target action responses (= correct responses) across the total number of sessions required to reach the learning criterion (80% correct responses in two consecutive sessions) for each action. To assess the effect of both action and group on the mean correct response rate of the learned actions, a generalized linear model (GLM) was fitted to predict mean response rate for all the actions. The fitted regression model was highly significant (R^2^ = 0.29, F (5, 49) = 5.428, p =0.00047) with group Test (β =0.25, p = 0.0007) and action ‘flap wings’ (β = -0.26, p = 0.02) correctly predicting the response rate. Thus, the subjects in the test condition showed significantly higher mean correct responses (p =0.00047) than the subjects in the test condition overall for the five target actions (see Figure 3), while for the action *‘flap wings’* the mean response rate was the least. After the first action was learnt, subjects in both groups initially continued to perform this very action when experimenters switched to the next novel command in the consecutive following sessions (a carry-over effect). But the subjects in the test condition switched to performing the second action faster than the subjects in the control condition (who learnt only two actions), which can be attributed as an effect of demonstration in the test group (see Supplementary Information Figure S3).

**Figure 3.**
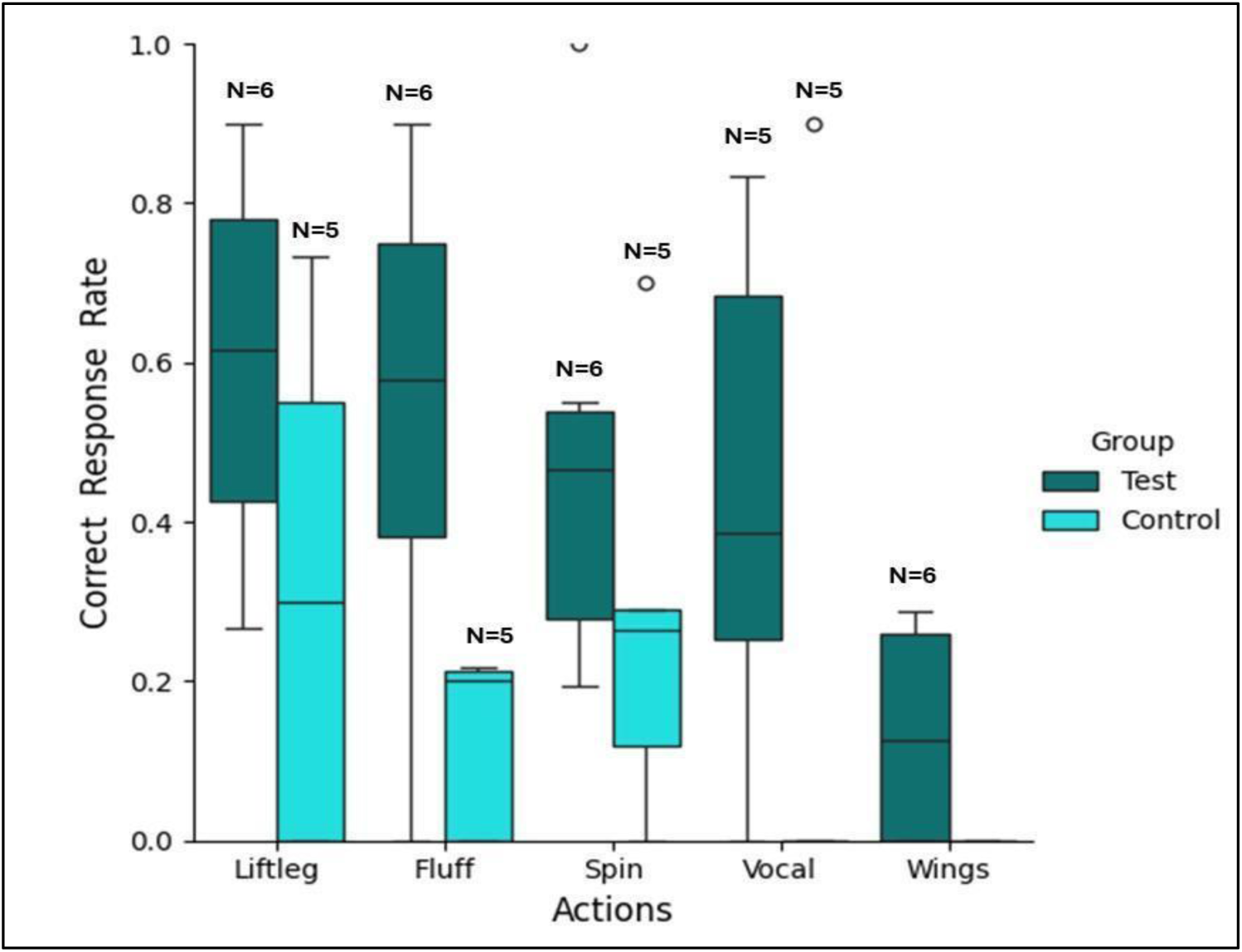
Mean correct response rate for each target action in the two experimental groups. The box plots show the mean response rate (y-axis) for each action (on the x-axis) for the test group (teal blue) and the control group (light blue). Error bars, mean± SEM

### Learning speed

The total number of sessions required to express each target action was considered as the learning speed for that action. In accordance with our predictions, the test group learned to express the target actions faster than the control group: a generalized linear model was fitted to test whether ‘group’ and ‘action’ significantly predicted this learning speed. The overall regression model was close to significance (R^2^ = 0.169, F (5, 30) = 2.42, p =0.058) with Test group correctly predicting the learning speed. The test group took significantly fewer number of sessions than the control group to learn to express the target actions upon command (β = -4.14, p = 0.0137). Figure 4 shows the mean learning speed of the two groups for each target action. Post-hoc analysis using a Wilcoxon-rank sum test showed that for the action *‘fluff’*, the five successful birds in the test group took significantly fewer sessions than the three successful birds in the control group (W=15, p=0.03) to learn to express this action. Such a significant difference was not observed for the other actions.

**Figure 4.**
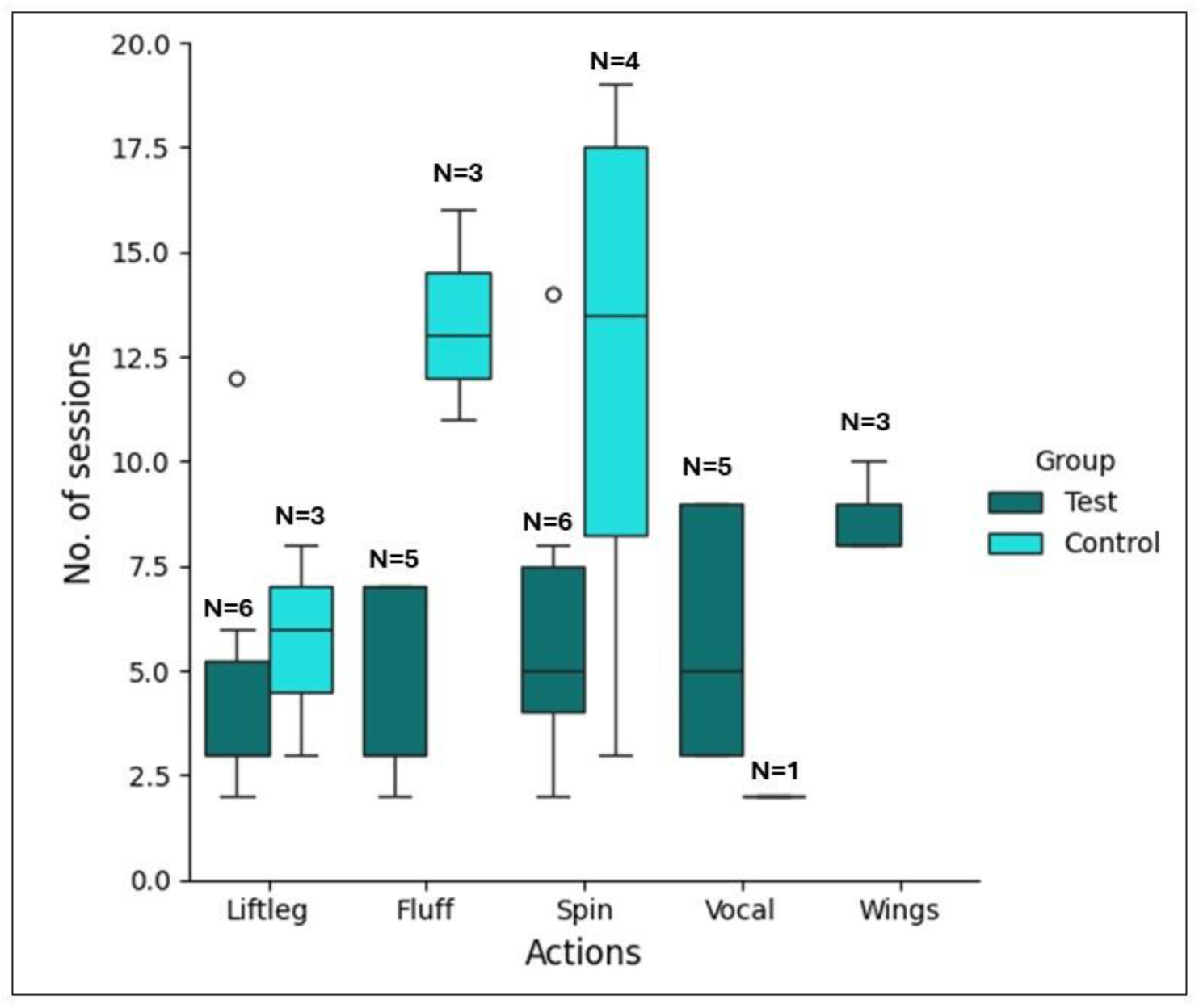
Mean number of sessions required to learn each target action. The box plots show the mean number of sessions (y-axis) required to learn each action (on the x-axis) for the test group (teal blue) and the control group (light blue). Error bars, mean± SEM

### Spontaneous imitation of target actions in the test group

In the test condition, a few cases of spontaneous imitation, which were not rewarded, occurred. In these cases, the subjects imitated the demonstrator in parallel to or shortly after the demonstration, i.e., within the 3-5 seconds gap before the hand command was given, prior to the first rewarded test trial (the production of the correct target action was only rewarded if it was produced in response to the hand command) of each action. Spontaneous imitation was observed with the actions *‘spin’* (all six test subjects produced *‘spin’* at least once, total events=21, Mean= 3.5, S.E=±1.0), *‘fluff’* (five subjects produced spontaneous *‘fluff’* at least once, total events= 10, Mean= 1.67, S.E=±0.71) and *‘vocal’* (two subjects produced *‘vocal’* at least once, total events= 4, Mean= 0.8, S.E=±0.58). For the actions *‘lift leg*’ and *‘wing’*, no spontaneous imitation occurred prior to the first rewarded test trial.

### Serial order of target actions learnt

To test whether the subjects would learn to express certain action orders differently (order effect), we fitted a generalized linear model with ‘order of the actions learnt’ as the response variable predicted by ‘action’ and ‘group’. The model was statistically significant (R^2^ = 0.36, F (5, 32) =5.29, p=0.001) with ‘action’ and ‘group’ (Group Test: β =0.7, p = 0.035) correctly predicting the order of the action learnt, showing that the subjects in the test group learnt particular actions first, followed by the (probably more difficult) actions later. The action *‘lift leg’* was usually the first action to be learnt, though the result was not statistically significant (β = -0.72, p = 0.14). *‘Flap wings’* was the last action (5^th^ in order) to be learnt for all subjects in the test group (β = 1.6, p = 0.008). No order effects were observed in the control group.

## DISCUSSION

Untrained, non-enculturated blue-throated macaws – a non-domesticated animal species – imitated intransitive actions in a third-party setting, providing the first evidence for third-party imitation outside humans. These birds learnt to produce five out of five improbable intransitive actions (improbability established via baseline data) by passively observing the communicative interaction between a trained conspecific and a human experimenter as a third party. In contrast, the birds in the control condition managed to learn only two of those actions on average and one action was not learnt at all. The finding that the macaws learned to express more target actions in response to human hand commands as well as learning it faster and more accurately in this third-party setting than when they did not observe the third-party interaction, corroborates that the macaws in the test group imitated the actions of the conspecific demonstrator in response to the human experimenter. Our findings strongly imply that the capacity to imitate as a third-party has convergently evolved in a non-mammalian taxon and suggests that macaws, and perhaps all parrots, may have evolved the ability to imitatively learn by observing the interactions of conspecifics without directly engaging with them, i.e., by third-party imitation.

The paradigm was demanding, considering that the parrots were not only required to imitate improbable intransitive actions, but also learn about the interactive context between the conspecific and the experimenter - the context in which the actions should be displayed (i.e., in response to a specific, arbitrary human gestural commands), in order to obtain a reward. Previously, only dogs have been tested and failed in a comparable third-party setting in which a demonstrator dog responded to verbal human commands novel to the subject (Tennie 2009). It is remarkable that dogs that (i) have life-long training of responding to human commands, (ii) exhibit a long history of domestication and (iii) can be trained to do motor imitation ^22,46^, fail to imitate in a third party setting spontaneously, whereas macaws, an undomesticated species, with naïve subjects that had never been trained to imitate or to perform any actions upon human instruction, succeeded. Moore ^16^ suggests that learning from third-party contexts requires an ability to recognise and imagine oneself in the third-party situation to correctly interpret a demonstration. This socio-cognitive skill is absent in one-year old human infants and develops from the second year of infancy ^12,13^. Thus, learning via third-party imitation represents a significant evolutionary step that has previously not been reported in other animal taxa ^17^.

In humans, learning social norms and traditions often require third-party imitation of community members in order to gain group affiliation and participate in group events ^47^. Most of these social conventions involve imitation of cognitively opaque arbitrary gestures and intransitive actions ^1^. Blue-throated macaws live in complex social groups, like most parrot species, with fission-fusion dynamics ^48^ which brings about frequent changes in group composition ^49^. This necessitates faster group synchronisation and integration into new social groups. Third-party imitative learning of intransitive actions from conspecifics may facilitate group integration, affiliation and social bonding and may eventually give rise to cultural conventions of coordinated movements or possibly even gestures in parrots. Future studies will show whether this hypothesis holds just for macaws or for other parrot species.

Unlike other studies investigating intransitive action imitation in non-human animals (in DAID or DATOD studies; see above), the macaws had not received any active training to imitate or ‘copy’ on command, nor were ostensive signals used to direct the subjects’ attention to the interaction. Vicarious reinforcement (subject observes the demonstrator getting rewards for performance) (50, 51) or observational conditioning (observer learns the relation between a stimulus and a reinforcer) (52), may have played a role, i.e., motivating the subjects to offer more actions that matched with the demonstrator’s after having observed a demonstrator receiving rewards for performing actions ^50^. However, correctly matching those demonstrated actions using the same effectors (body parts) in response to specific hand commands within a short period of time requires more than just motivational incentives, making third-party imitation the more parsimonious explanation.

Against our predictions, the control subjects succeeded in acquiring the correct response to some of the commands (on average 2.2). A closer examination of what actions they learnt suggests that the birds did not just learn by trial-and-error. Two of the gestural commands may have involved some unintended cueing that led to the acquisition of those actions by the control group. For example, *‘spin’*, which was learnt by four control individuals seems to have been cued by the spin-like movement of the experimenter’s hand. The experimenter turned her two outstretched fingers in the anticlockwise direction which might have induced the control subjects to follow the finger movement and rotate in the same direction. The test subjects however, rotated in the opposite direction to the movement of the fingers (see Supplementary Informations) copying the anticlockwise rotation of the demonstrator instead. The test group therefore imitated the demonstrator rather than being influenced by cues conveyed by the gestural commands. *‘Lift leg’*, which was acquired by three individuals in the control group (and all six individuals in the test group), may have been learnt via a potential correspondence of a lifted foot to the lifted index finger ^51^ which constituted the gestural command. The remaining three commands however could not possibly have provided any potential cue towards the target action and were either not learnt at all by the control group (in the case of *‘flap wings’*) or by fewer subjects than in the test group (concerning ‘*fluff’* and *‘vocal’*). The action ‘flap wings’ was also learnt last in the order by the three test subjects who acquired it, implying the improbability of the behaviour as evidenced by the very low occurrence rate in the baseline observations. These actions linked with completely arbitrary hand commands learnt by the test group but mostly not learnt by the control group provide robust evidence for third-party imitation. The finding that some control subjects successfully produced correct responses to novel human gestures, even though they had never been trained to produce behaviours in interaction with a human experimenter, is very interesting in itself and suggests that the species has a very strong communicative tendency that even transcends species boundaries.

For some actions (‘*spin’, ‘fluff’, ‘vocal’*) few instances of spontaneous imitation were observed in the test group concurrently with demonstrations before receiving the gestural command. Such instances occurred even without any prior reinforcement of that action. Those unrewarded imitation events show spontaneous, direct imitation and imply that parrots may have naturally evolved a tendency to imitate intransitive actions from conspecifics. In the large majority of cases however, the test group parrots clearly showed the action only in response to the gestural command, thus replicating the appropriate response witnessed as in a third-party context.

Overall, our study provides the first evidence for spontaneous imitative learning from a third-party setting in a parrot, an ability previously recorded only in humans and, concurrently, the first evidence for third-party imitation of intransitive actions in a parrot. Our findings show that blue-throated macaws learn imitatively from observing conspecific interactions, suggesting that third-party imitation of intransitive actions may be ubiquitous in parrots. This may facilitate faster transmission of social information and group synchronisation that could provide adaptive benefits to a highly social species ^52^. It also raises the possibility that macaws, and perhaps other parrots, may transmit and use gestures and bodily movements in the wild, maybe to a degree (not tested here) that may give raise to long-lasting cultures. Our study opens up a new line of inquiry into third party imitation, and the imitation of intransitive actions, its evolution, and its potential importance for group cohesion and cultural evolution in a non-primate taxon: parrots.

## MATERIALS AND METHODS

### Subjects and housing conditions

Subjects were 13 (experimental subjects N=11, demonstrators N=2) hand-raised but early-on socialized and group-living captive blue-throated macaws (for details on rearing, age and sex refer to Supplementary Information and **Table S2**). Prior to the study, they were kept at the Max Planck Comparative Cognition Research Station facility at Loro Parque Fundación, Tenerife, Spain. All subjects were group-housed in 3 adjacent semi-outdoor aviaries contiguous with the lab facility. All birds had *ad libitum* access to water and mineral blocks and were fed twice daily (for more details on housing and diet, please refer to Supplementary Information). Besides the two pre-assigned trained demonstrators, 11 subjects were divided into two groups (balancing for age and sex): i) a test group that was tested in a third-party setting with demonstrator (n=6) and ii) a control group (n=5) tested without a demonstrator in the first round and with a ‘social facilitator’ (as social enhancement control) in the second round (see below). The two demonstrators (8-year-old males) were behaviourally trained. They could perform 5 intransitive motor actions upon specific gestural commands (hand signals) from the experimenter (see Supplementary Information Figure S1). In contrast, the test and control subjects were completely naïve individuals that had not participated in any prior cognitive study previously and were only habituated to human presence, stay on a perch in the experimental room and receive food from human hands. They had not been trained to perform any behaviour on command or to copy actions of others. All subjects participated in the experiment voluntarily.

### Intransitive actions used in the experiment

Five arbitrarily chosen intransitive motor actions, namely i) *‘lift leg’* ii) *‘spin’* iii) *‘fluff’* iv) *‘vocal’* and v*) ‘flap wings’* were used for the experiment. The detailed topography of each demonstrated action with the associated commands is described in Supplementary Information Table S1 and illustrated in Supplementary Figure S1. The order in which the five different actions and associated commands were introduced was randomized across subjects for both test and control group.

### Baseline: Natural occurrence rate of the target actions

Behavioural observations were conducted throughout 16 days before the experiment commenced. We monitored each of the 11 subjects for 15 minutes (one control subject was not available) per day throughout 16 days, resulting in 44h of observation in total. During each one-minute observation, it was noted if any of the five focal behaviours were shown by each subject. The subjects were observed by the experimenter at different times of the day when they were not feeding, following some criteria (see Supplementary Information). The occurrences of each of those five actions from all the subjects in 44 hours were summed to obtain cumulative scores (Supplementary Information). The rate of occurrence within a 12 secs timeframe was then calculated as *Rate = (Cumulative score X 12)/44 X 60 X 60*. We set a post-hoc criteria for the rate of occurrence to infer which behaviours are more frequently observed in the natural repertoire-i) Below 10%-infrequent, ii) Below 5%-improbable and iii) Below 1%-very improbable.

### Experimental setup

An illustration of the experimental set-up is shown in Figure 1. Testing took place in two adjacent indoor testing chambers, separated from each other with a transparent plexiglass window and equipped with lamps covering the birds’ full range of visible light (details in Supplementary Information), In each room, a standing perch (height: 1.13m) was placed at a 1m distance on either side of the plexiglass window. The demonstrator was placed on the perch to the left side of the glass while the observer was placed on the right side. Experimenter 1 stood in chamber 1 facing the demonstrator while experimenter 2 stood in the neighbouring chamber facing the subject. Both experimenters wore reflective sunglasses to avoid any unconscious cueing. The setup for the control sessions was kept the same as the tests with the control subject sitting on the perch in front of experimenter 2. The adjacent chamber was empty for the first round of testing in the control condition. From the second round, a conspecific bird (social facilitator)^50^ was placed on the perch accompanied by experimenter 1 – as a control for social enhancement ^53^. We recorded the sessions on cctv cameras mounted on the walls of the chamber and a Sony HDR-CX240E video camera was additionally used. Testing took place from October 2021 to August 2022 between 10:30 and 14:00 regularly every week.

### General procedure

#### Reward contingency

The demonstrator was rewarded with a small piece of walnut for each correct demonstration in clear view of the subjects. No other actions of the demonstrator except the requested action during a trial was rewarded. The social facilitator in the control sessions was fed sunflower seeds every 2-3 minutes. The subjects in both conditions were rewarded with small pieces of walnuts for the target actions during the test trials. Any action that was not the target action, or that was performed outside of the 12 (±2) sec interval (set *a priori*) following the gestural command given by experimenter 2 (i.e., the actual duration of the test trial, see below) was not rewarded.

#### Test condition

In each test session only one of the five actions was demonstrated and tested. At the beginning of each test session, the demonstrator and the subject were placed next to each other on the two standing perches Both birds could clearly see both experimenters. Test sessions consisted of the 10 trials preceded by three demonstrations each, hence 30 demonstrations and 10 trials in total. Experimenter 1 gave the gestural command requesting the demonstrator to perform a particular action (e.g., *‘lift leg’*)’. As soon as the demonstrator performed the action, she pressed a clicker and rewarded the bird with a sunflower seed in full view of the subject (for a description of the visual field of parrots see Demery et al. 2011)^54^.

The three demonstrations were carried out in a row by experimenter 1 interacting with the demonstrator, then, after a gap of 3-5 seconds, experimenter 2 started the test trial. She gave the same gestural command to the subject for 12 (±2) seconds continuously, which experimenter 1 had given throughout the demonstrations (see Supplementary Videos S1-5). If the subject performed the target action within this period of time, experimenter 2 pressed the clicker immediately and rewarded the subject with a walnut piece. If there was no response from the subject or a different action was performed, there was no reward, the response was noted as zero, and experimenter 1 continued with the subsequent trial.

##### a) Round 1 of testing

To avoid frustration of the subjects in case of continuous failure, we adopted a ‘rolling’ method for demonstration of the actions in two consecutive rounds. In the first round, the first action (as determined from the individually randomized order provided in supplementary Table S2) was demonstrated throughout a minimum of four sessions. In absence of any correct matching response from the subject within those four consecutive sessions, the 2^nd^ action was demonstrated from the next session throughout at least four sessions, and the process continued for the following actions. If there was any correct response from the subject within those four sessions of a particular action, the testing continued until the subject reached the learning criterion of 80% correct responses (8 out of 10 correct trials) in two consecutive sessions. After all the 5 actions had been demonstrated in at least 4 sessions, round 2 commenced.

##### b) Round 2 of testing

Round 2 followed the same protocol as round 1, but now the minimum number of test sessions for an action was increased to 7 in a row. All actions that the subjects had failed to learn in round 1, were repeated in round 2 (see Supplementary Information Table S2). If a subject failed to respond correctly within those seven sessions, it was considered not to have learnt the action. If there was any target matching response within these seven sessions, the testing continued with that action until the subject had reached the 80% learning criterion in two consecutive sessions as in round 1.

### Control condition

The control sessions consisted of the same rolling method of two rounds employing the same number of sessions as the test condition. The first round was conducted without a conspecific present in the adjacent test chamber. However, in order to control for possible effects of social enhancement, a 2^nd^ bird (social facilitator) ^53^ was placed in the adjacent chamber from round 2 in the presence of experimenter 1. Experimenter 1 stood passively in front of the social facilitator, not giving any commands or cues. Occasionally, experimenter 1 would give sunflower seeds to the social facilitator to prevent it from begging using vocalisations or doing any other action. Experimenter 2 would give a hand signal to the subject for 12 (±2) seconds continuously for the bird to respond (see supplementary videos S6-7). If the subject produced the target action, a clicker was pressed immediately followed by rewarding of the subject with a walnut piece, otherwise the next trial ensued after an inter-trial gap of 3-5 seconds. There were 10 trials in each session and the criteria for successful learning of a behaviour or failure were kept identical to the test condition. The social facilitator was rewarded only with sunflower seeds which was of lower value than walnut, to ensure the control subjects would not socially learn to remain passive like the social facilitator.

### Analyses

The videos were coded by two experimenters (who participated in the experiment) using Solomon Coder (version Beta 19.08.02, 2019 by András Péter). Production of target action within 12 (±2) sec after the hand signals in each session were coded as successful trials. Absence of any response or non-target responses were coded as failed trials. The total number of sessions faced in round 1 and round 2 to learn each behaviour was taken as the learning speed. For example, if a bird did not express any target action response in the four sessions of the1^st^ round but learnt to express the target action in the 2^nd^ round taking 13 sessions, we took the total session number as 17. All statistical analyses (Wilcoxon-rank sum test and GLM) were performed in RStudio (R Core Team 2019, RStudio version R.4.1.2).

## ETHICAL STANDARDS

All applicable international, national, and/or institutional guidelines for the care and use of animals were followed. Under the German Animal Welfare Act of 25th May 1998, Section V, Article 7 and the Spanish Animal Welfare Act 32/2007 of 7th November 2007, Preliminary Title, Article 3, the study was not classified as an animal experiment and did not require any approval from a relevant body. The experiments did not require an application to the Animal Ethics Committee of neither Germany or Spain, as animals participated voluntarily in the experiments and were not affected by them in any way. This article does not report any studies with human participants performed by any of the authors. The ARRIVE guidelines for the reporting of animal experiments were followed.

## DATA AVAILABILITY

Raw and analysed data with R codes have been deposited at Figshare and is publicly available at DOI: 10.6084/m9.figshare.26819929

### Code

This paper reports R codes for known statistical analysis like GLM and Wilcoxon-rank sum tests.

### Materials availability

Detailed description of the materials is listed in the Methods and Supplementary section. Any further information is available from the corresponding authors E.H and A.M.P.B.

## ACKNOWLEDGEMENTS AND FUNDING

We thank Loro Parque and its president Mr. Wolfgang Kiessling for their generous support, the access to the birds and the research facilities. We thank Loro Parque Foundation, its president Mr Christoph Kiessling, and staff, especially the animal caretakers and the veterinary department, for their constant support. We thank Caterina Bonizzato for the illustrations of the hand commands, Deepshika Arun, Francesca Castellano for assisting with the data collection. Esha Haldar received funding from DAAD Graduate School Scholarship Program and Animal Minds Project e.V.. Sara Torres Ortiz received funding from Animal Minds Project e.V.

## AUTHOR CONTRIBUTIONS

E.H and A.M.P.B conceived the study. E.H. and A.M.P.B designed and coordinated the study. E.H and S.T.O, J.V, M.V, A.H.S collected the data, E.H, J.V and M.V analysed the data, E.H wrote the manuscript. C.T and A.M.P.B edited the manuscript. All authors gave final approval for publication and agree to be held accountable for the work performed therein.

## COMPETING INTERESTS

The authors declare no competing interests.

## SUPPLEMENTAL INFORMATION

Supplementary Information SI

Video S1. Exemplary video of a *‘spin’* test trial

Video S2. Exemplary video of a *‘lift leg’* test trial

Video S3. Exemplary video of a *‘fluff’* test trial

Video S4. Exemplary video of a *‘vocal’* test trial

Video S5. Exemplary video of a *‘flap wings’* test trial

Video S6. Exemplary video of a control trial *(‘vocal’*)

Video S7. Exemplary video of a control trial *(‘lift leg’*) with social facilitator present

## Notes

### Competing Interest Statement

The authors have declared no competing interest.

## REFERENCES

1. Gergely, G. & Csibra, G. Sylvia’s Recipe: The Role of Imitation and Pedagogy in the Transmission of Cultural Knowledge. in Roots of Human Sociality (eds. Enfield, N. J. & Levinson, S. C.) 229–255 (Routledge, 2020). doi:10.4324/9781003135517-11.

2. Jagiello, R., Heyes, C. & Whitehouse, H. Tradition and invention: The bifocal stance theory of cultural evolution. Behav Brain Sci 45, e249 (2022).

3. Matsumoto, D. & Hwang, H. C. Cultural Similarities and Differences in Emblematic Gestures. J Nonverbal Behav 37, 1–27 (2013).

4. Takada, A. Education and Learning During Social Situations Among the Central Kalahari San. in Social Learning and Innovation in Contemporary Hunter-Gatherers (eds. Terashima, H. & Hewlett, B. S.) 97–111 (Springer Japan, Tokyo, 2016). doi:10.1007/978-4-431-55997-9_8.

5. Csibra, G. & Gergely, G. Social learning and social cognition: The case for pedagogy. in *Processes of Change* in Brain and Cognitive Development (eds. Munakata, Y. & Johnson, M. H.) 249–274 (Oxford University PressOxford, 2006). doi:10.1093/oso/9780198568742.003.0011.

6. Boyette, A. H. & Hewlett, B. S. Teaching in Hunter-Gatherers. Rev.Phil.Psych. 9, 771–797 (2018).

7. Kline, M. A. How to learn about teaching: An evolutionary framework for the study of teaching behavior in humans and other animals. Behav Brain Sci 38, e31 (2015).

8. Rogoff, B., Paradise, R., Arauz, R. M., Correa-Chávez, M. & Angelillo, C. Firsthand Learning Through Intent Participation. Annu. Rev. Psychol. 54, 175–203 (2003).

9. Salali, G. D. et al. Development of social learning and play in BaYaka hunter-gatherers of Congo. Sci Rep 9, 11080 (2019).

10. Gaskins, S. & Paradise, R. Chapter five: Learning through observation in daily life. The anthropology of learning in childhood, 85, 85–110. in The anthropology of learning in childhood 85–110 (2010).

11. Paradise, R. & De Haan, M. Responsibility and Reciprocity: Social Organization of Mazahua Learning Practices. Anthropology & Edu Quarterly 40, 187–204 (2009).

12. Herold, K. H. & Akhtar, N. Imitative learning from a third-party interaction: Relations with self-recognition and perspective taking. Journal of Experimental Child Psychology 101, 114–123 (2008).

13. Matheson, H., Moore, C. & Akhtar, N. The development of social learning in interactive and observational contexts. Journal of Experimental Child Psychology 114, 161–172 (2013).

14. Floor, P. & Akhtar, N. Can 18-Month-Old Infants Learn Words by Listening In on Conversations? Infancy 9, 327–339 (2006).

15. Stenberg, G. Infant imitation in a third-party context. IS 21, 387–411 (2020).

16. Moore, C. Understanding self and others in the second year. Socioemotional development in the toddler years: Transitions and transformations, 43–65. in Socioemotional Development in the Toddler Years: Transitions and Transformations 43–65.

17. Tennie, C. et al. Dogs, *Canis familiaris*, fail to copy intransitive actions in third-party contextual imitation tasks. Animal Behaviour 77, 1491–1499 (2009).

18. Heyes, C. & Saggerson, A. Testing for imitative and nonimitative social learning in the budgerigar using a two-object/two-action test. Animal Behaviour 64, 851–859 (2002).

19. Voelkl, B. & Huber, L. True imitation in marmosets. Animal Behaviour 60, 195–202 (2000).

20. Akins, C. K., Klein, E. D. & Zentall, T. R. Imitative learning in Japanese quail (*Coturnix japonica*) using the bidirectional control procedure. Animal Learning & Behavior 30, 275–281 (2002).

21. Zentall, T. R., Sutton, J. E. & Sherburne, L. M. True Imitative Learning in Pigeons. Psychol Sci 7, 343–346 (1996).

22. Topál, J., Byrne, R. W., Miklósi, Á. & Csányi, V. Reproducing human actions and action sequences: “Do as I Do!” in a dog. Anim Cogn 9, 355–367 (2006).

23. Custance, D. M., Whiten, A. & Bard, K. A. Can Young Chimpanzees (Pan troglodytes) Imitate Arbitrary Actions? Hayes & Hayes (1952) Revisited. Behaviour 132, 837–859 (1995).

24. Call, J. BODY IMITATION IN AN ENCULTURATED ORANGUTAN ( *PONGO PYGMAEUS*). Cybernetics and Systems 32, 97–119 (2001).

25. Abramson, J. Z. et al. Contextual imitation of intransitive body actions in a Beluga whale (Delphinapterus leucas): A “do as other does” study. PLoS ONE 12, e0178906 (2017).

26. Abramson, J. Z., Hernández-Lloreda, V., Call, J. & Colmenares, F. Experimental evidence for action imitation in killer whales (Orcinus orca). Anim Cogn 16, 11–22 (2013).

27. Jaakkola, K., Guarino, E. & Rodriguez, M. Blindfolded Imitation in a Bottlenose Dolphin (*Tursiops truncatus*). International Journal of Comparative Psychology 23, (2010).

28. Range, F., Viranyi, Z. & Huber, L. Selective Imitation in Domestic Dogs. Current Biology 17, 868–872 (2007).

29. Tomasello, M., Call, J., Nagell, K., Olguin, R. & Carpenter, M. The learning and use of gestural signals by young chimpanzees: A trans-generational study. Primates 35, 137–154 (1994).

30. Tomasello, M. & Zuberbühler, K. Primate Vocal and Gestural Communication. in The Cognitive Animal (eds. Bekoff, M., Allen, C. & Burghardt, G. M.) 293–300 (The MIT Press, 2002). doi:10.7551/mitpress/1885.003.0041.

31. Byrne, R. W. et al. Great ape gestures: intentional communication with a rich set of innate signals. Anim Cogn 20, 755–769 (2017).

32. Tomasello, M. & Call, J. Thirty years of great ape gestures. Anim Cogn 22, 461–469 (2019).

33. Halina, M., Rossano, F. & Tomasello, M. The ontogenetic ritualization of bonobo gestures. Anim Cogn 16, 653–666 (2013).

34. Moore, B. R. Avian Movement Imitation and a New Form of Mimicry: Tracing the Evolution of a Complex Form of Learning. Behav 122, 231–263 (1992).

35. Todt, D. Social Learning of Vocal Patterns and Modes of their Application in Grey Parrots (*Psittacus erithacus* ) ^1, 2, 3^. Zeitschrift für Tierpsychologie 39, 178–188 (1975).

36. Pepperberg, I. M. A Review of the Model/Rival (M/R) Technique for Training Interspecies Communication and Its Use in Behavioral Research. Animals 11, 2479 (2021).

37. Chakraborty, M. et al. Core and Shell Song Systems Unique to the Parrot Brain. PLoS ONE 10, e0118496 (2015).

38. Carouso-Peck, S., Goldstein, M. H. & Fitch, W. T. The many functions of vocal learning. Phil. Trans. R. Soc. B 376, 20200235 (2021).

39. Benedict, L., Charles, A., Brockington, A. & Dahlin, C. R. A survey of vocal mimicry in companion parrots. Sci Rep 12, 20271 (2022).

40. Auersperg, A. M. I. et al. Social transmission of tool use and tool manufacture in Goffin cockatoos ( *Cacatua goffini* ). Proc. R. Soc. B. 281, 20140972 (2014).

41. O’Hara, M. et al. Wild Goffin’s cockatoos flexibly manufacture and use tool sets. Current Biology 31, 4512–4520.e6 (2021).

42. Aplin, L. M. et al. Experimentally induced innovations lead to persistent culture via conformity in wild birds. Nature 518, 538–541 (2015).

43. Klump, B. C. et al. Innovation and geographic spread of a complex foraging culture in an urban parrot. Science 373, 456–460 (2021).

44. Wright, T. F. & Dahlin, C. R. Vocal dialects in parrots: patterns and processes of cultural evolution. Emu - Austral Ornithology 118, 50–66 (2018).

45. Mui, R., Haselgrove, M., Pearce, J. & Heyes, C. Automatic imitation in budgerigars. Proc. R. Soc. B. 275, 2547–2553 (2008).

46. Fugazza, C., Petro, E., Miklósi, Á. & Pogány, Á. Social learning of goal-directed actions in dogs (*Canis familiaris*): Imitation or emulation? Journal of Comparative Psychology 133, 244–251 (2019).

47. Van Schaik, C. P. Social learning and culture in animals. in Animal Behaviour: Evolution and Mechanisms (ed. Kappeler, P.) 623–653 (Springer Berlin Heidelberg, Berlin, Heidelberg, 2010). doi:10.1007/978-3-642-02624-9_20.

48. Hesse, A. J. & Duffield, G. E. The status and conservation of the Blue-Throated Macaw *Ara glaucogularis*. Bird Conservation International 10, 255–275 (2000).

49. Manual of Parrot Behavior. (Wiley, 2006). doi:10.1002/9780470344651.

50. Zentall, T. R. Imitation: definitions, evidence, and mechanisms. Anim Cogn 9, 335–353 (2006).

51. Nehaniv, C. L. & Dautenhahn, K. The Correspondence Problem. in Imitation in Animals and Artifacts (eds. Dautenhahn, K. & Nehaniv, C. L.) 41–62 (The MIT Press, 2002). doi:10.7551/mitpress/3676.003.0003.

52. Caldwell, C. & Whiten, A. Evolutionary perspectives on imitation: is a comparative psychology of social learning possible? Animal Cognition 5, 193–208 (2002).

53. Franz, M. & Matthews, L. J. Social enhancement can create adaptive, arbitrary and maladaptive cultural traditions. Proc. R. Soc. B. 277, 3363–3372 (2010).

54. Demery, Z. P., Chappell, J. & Martin, G. R. Vision, touch and object manipulation in Senegal parrots *Poicephalus senegalus*. Proc. R. Soc. B. 278, 3687–3693 (2011).

