## Supplementary Information for "Third-party imitation is not restricted to humans"

### Supporting Information Text

#### Supplementary Methods

*Rearing conditions of the subjects*

At Loro Parque, many parrot chicks (including all our subjects) are hand-raised every year by professionals. However, as soon as they can perch, they are put together with conspecifics and have no human contact except for the handfeeding and as soon as they feed independently (ca 6 weeks old), they are socialized and group housed in large aviaries with conspecifics. The subjects were brought to the Max-Planck comparative cognition research station shortly before the experiment commenced (see Table S2 for the ages of the different individuals). After they had got accustomed to their new environment and new home aviaries, they were observed throughout 16 days in those aviaries for the baseline data collection. In parallel the habituation to the experimental rooms and to the experimenters took place. Before the experiment commenced the subjects had been habituated to basic handling and clickers, remaining on the perch in the experimental chamber in the presence of a human experimenter and to receiving rewards from human hands. Otherwise, they had received no further training.

*Housing conditions*

All subjects were group-housed in 3 adjacent semi-outdoor aviaries contiguous with the lab facility. The aviaries measured approximately 1.80 × 3.40 × 3 m (width × length × height) with interconnected windows (1m x 1m) which remained closed throughout the experiment to avoid social conflicts between certain individuals. All birds had 24-hour access to the outside aviary, allowing them to follow a natural light cycle. Half of the aviary was outdoors so the birds were exposed to natural weather conditions. The other half was covered and lit with Arcadia Zoo Bars (Arcadia 54W freshwater Pro and Arcadia 54W D3 Reptile Lamp) that automatically followed the natural daylight regime. Outdoor temperatures during the research period fluctuated between 20 to 26 degrees Celsius during the day and 15 to 21 degrees Celsius during the night with interspersed periods of light raining. All birds had ad libitum access to water and mineral blocks. They were fed with fresh fruits, vegetables twice a day along with a seed mix only in the afternoon. Their daily nut ration was provided to them throughout the testing, training or enrichment sessions. To transport the subjects to the testing rooms, the aviaries were connected with mobile (1m x 1m x 1m) feeding cages which could be wheeled with ease with the birds inside. The birds were completely used to this procedure and entered the cages to be transported readily and voluntarily.

*Experimental setup*

Testing took place in two adjacent indoor testing chambers, separated from each other with a transparent plexiglass window and equipped with lamps covering the birds' full range of visible light (Arcadia Zoo Bars). Measurement of each testing chamber was 2.5 m × 1.5 m × 1.5 m (height × width × length). The chambers were separated from each other with a transparent plexiglass window. of 1.0 m x 1.0 m with an opening in the lower part of the glass so that the parrots and the experimenters could clearly see and hear each other. The experiments could be observed by the Loro Parque zoo visitors through a one-way window such that the birds could not see or hear anything from outside the wall.

*Gestural commands and associated target actions*

Following the 3 demonstrations of the third party (three times command given and target action shown in response followed by clicker!) the same gestural commands were given to the test and control subjects throughout 12(+-2) seconds time period in each test trial. If the subject produced a correct response within that time period, it was immediately followed by a clicker and a consecutive reward.

#### Supplementary Results

*Baseline: Natural occurrence rate of the target actions*

Except for self-defending from a conspecific attack, the action '*lift leg*' was not observed. Only transitive versions of *'lift leg'* which we did not count, were observed, such as scratching of head or eyes with the leg, tucking in of one leg while sleeping or resting and nibbling of the ring or digits while lifting the foot. Hence, '*lift leg'* as considered as the target response, never occurred. Defending from attack occurred 4x, scratching 84x, resting 86x, nibbling 52x. The most frequently occurring action that was recorded was expectedly *'vocal'* (657x), followed by *'fluff'* (68x). Flapping wings while sitting was observed 6x while full rotation on a perch as approximation to *'spin'* action was observed only once. The rate of occurrence at 15 secs timeframe was calculated as *Rate=(Cumulative score X 12)/44 X 60 X 60*.

*Action topography differences between the two experimental groups*

(i) *'Lift leg'*

There was no difference in the form of the action between the two groups. The successful subjects lifted either of their feet and held it up in the air near the chest with four digits splayed out. There was no preference for a particular foot in either of the groups.

(ii) *'Spin'*

An interesting difference was observed between the test and the control group for the action spin. Four out of six (66%) individuals in the test group preferred to spin in clockwise direction which was mirroring the direction of the demonstrator's movement, i.e., rotating in the opposite direction. In contrast, the four control subjects who learnt the action, span in the anticlockwise direction exactly like the trained demonstrator, i.e., the same direction as the circular movement of the experimenter's forehand. The explanation for this difference lies in conceptualising '*spin*' not as one, but a multi-step action that the test subjects copied from the demonstrator. This involves, starting from facing the experimenter, turning towards the window separating the rooms, facing opposite to the experimenter, then away from the other bird and lastly back to the experimenter (see Figure S2).

(iii) *'Fluff'*

*'Fluff'* was characterized by a head shake with the feathers around the head and neck standing up as if aroused. There was no visible difference between the performance of the two groups in this action.

(iv) *Vocal*

As we considered any call or vocalisation as the target action for 'vocal', we did not analyse the spectrograms of the test subjects for similarity with the demonstrator or controls.

(v) '*Flap wings'*

*'Flap wings'* was the last action the subjects in the test group happened to learn. None of the controls learnt the action wings confirming the improbability of the action to occur naturally. One control subject repeatedly made characteristic begging movements with its wings accompanied by vocalisations. The topography of the begging movement did not match the target action and therefore was not rewarded. Another test subject, the youngest half year old individual, also showed begging movement with its wings, which differed

noticeably from the demonstration. This was also considered an incorrect response and was not rewarded. Half of the subjects in the test group were able to learn the action matching closely with the demonstrator.

*Effect of age*

We tested whether age could affect the social learning ability in this species as well as explain the non-responsiveness of the single individual in the test group which was excluded from the previous analysis. To test the effect of age on the imitative learning ability of the individuals, we fitted a linear regression model with age predicting the number of actions learnt by the test group subjects by observing a conspecific. When we included the individual who did not learn any single behaviour in our model, the prediction was stronger with number of actions learnt significantly decreasing with age ($F_{(1,5)} = 8.188$, $\beta = -0.24$, $R^2$ of 0.545, $p = 0.035$). There was no difference between the young and the adults of the control group in their trial-and-error learning ($\beta = -0.04$, $p = 0.51$). **Figure S4** shows that young macaws are more prone to social learning of arbitrary actions from a conspecific demonstrator than the adults.

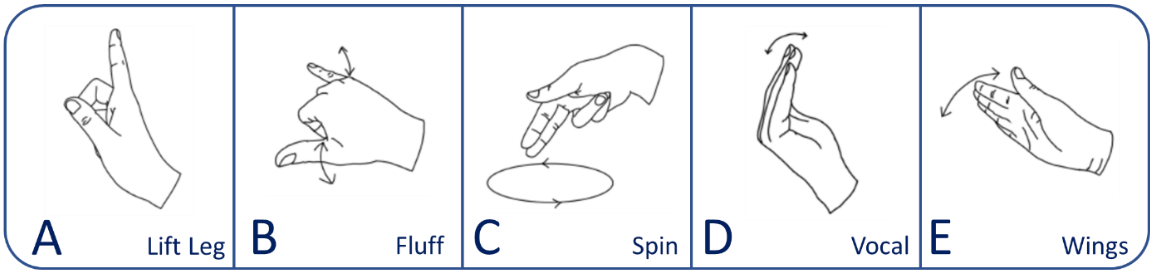

Fig. S1. Illustration of the gestural commands associated with the target actions

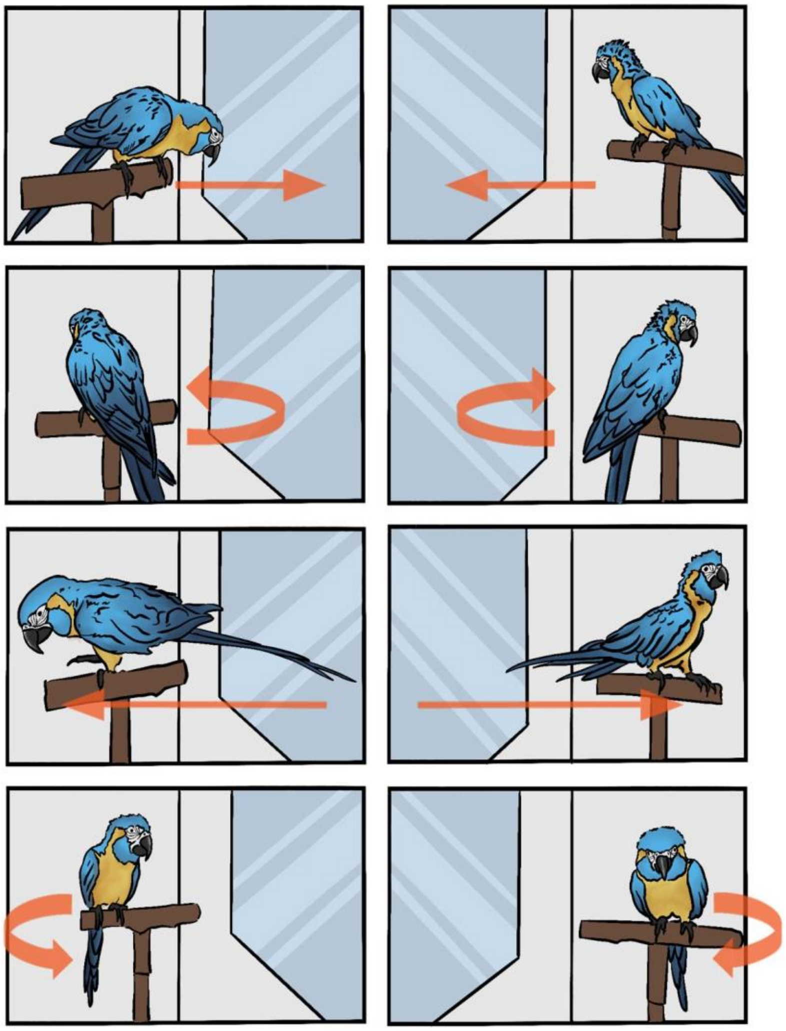

Fig. S2. ‘Spin’ target action of a test subject. The demonstrator is on the left panel and the test subject is on the right panel. The movement of the test subject during a ‘spin’ is shown in four consecutive steps.

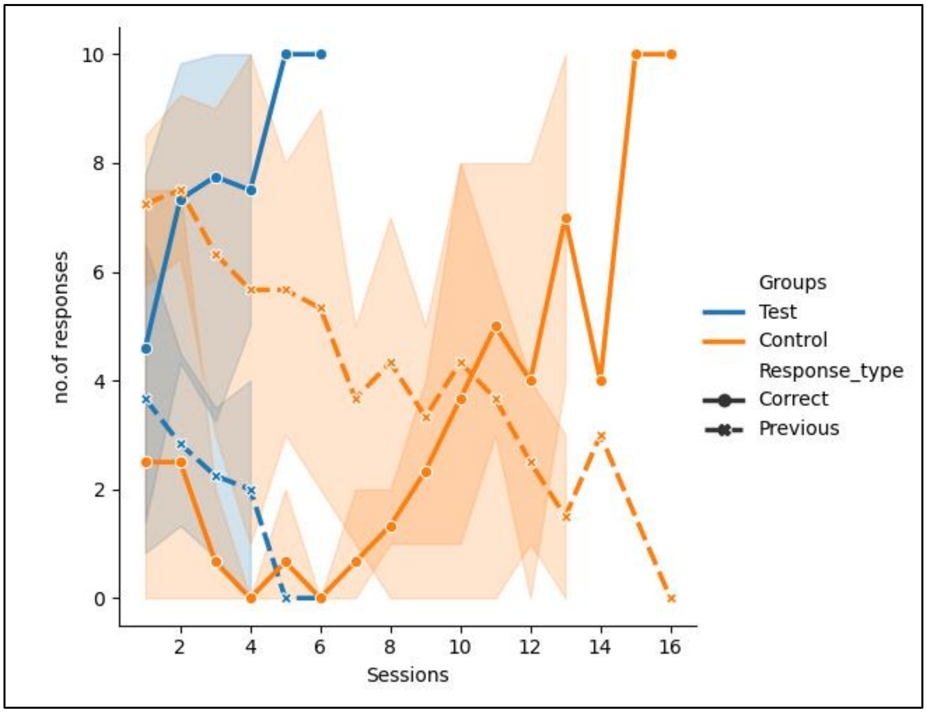

Fig. S3. Carry-over effect. Mean ‘correct responses’ of current action and mean of ‘previous action’ responses (first action learnt) are plotted across total number of sessions for the two experimental groups. At (0,0) subjects had learnt their preceding (first) action, followed by different response types (correct and previous) in the following sessions leading to learning of the current action denoted by correct responses above 80%. We did not consider the action switch for one control subject (Dr. Strange) who was intermittently absent from our research station and may have had some close human contact in the veterinary station. The subject had not produced any response in the first round of testing, but after the intermittent absence, it learnt two actions in the second round. This quick uptake of the behaviours may have been influenced by external factors, and hence we did not consider him for the action switching analysis.

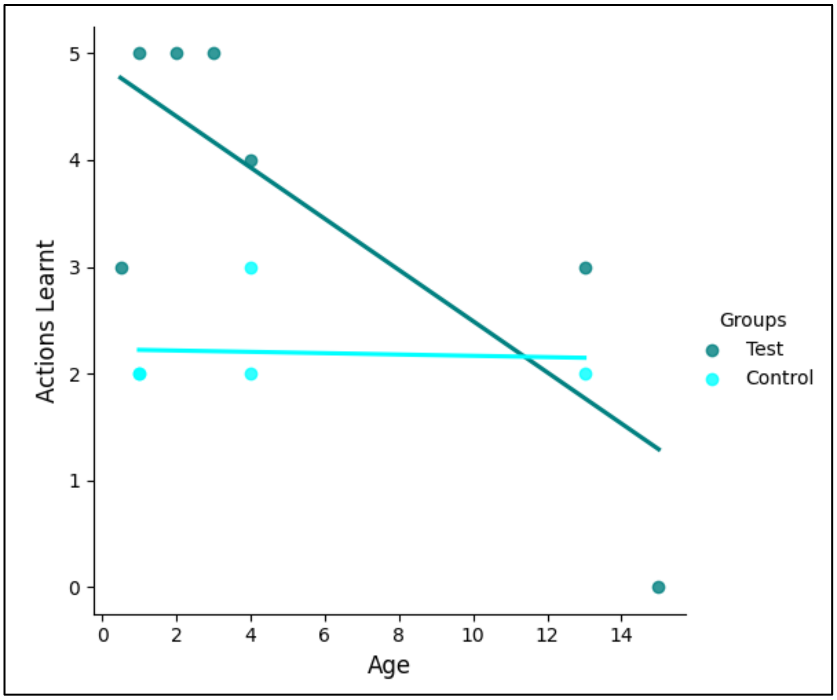

Fig. S4. **Scatter plot with regression lines showing the number of target actions learnt as function of age of macaws in the test (teal blue) and control (light blue) condition.** The number of actions learnt decreased significantly with age in the test condition when the subject who did not learn any action was included (linear regression, ($F_{1,5}$=8.188, β= -0.24, $R^2$ of 0.545, p= 0.035). The linear regression was not significant for the control condition ((β =-0.04, p =0.51).

**Table S1.** Description of the target actions and the gestural commands used in the experiment

| Target actions | Topography of demonstrations | Gestural Commands | Criteria for correct response |
|---|---|---|---|
| 1. Lift leg | Demonstrator lifts the right foot up with splayed out digits and holds it in the air near the chest. | Experimenter holds the pointed index finger of her right arm at a chest level of the subject keeping an optimum distance from the subject. | Any foot can lifted up and held in the air for some time, close to the chest, with digits splayed out (i.e. without grasping any object) |
| 2. Spin | Demonstrator makes a 360 degree turn on the perch in an anticlockwise direction. | Experimenter lifts her right arm up, little above the eye level of the subject and uses two pointed fingers to make repeated anticlockwise round motions. | Subject has to complete 360 degree rotation on the perch in any direction (clock or anticlockwise). |
| 3. Fluff | Demonstrator rapidly shakes the head. | Experimenter folds the three middle fingers and splays out the thumb and little finger of right arm. She holds the right | Shaking of head is only considered, not the whole body. |

| | | | |
|---|---|---|---|
| | | arm up at the chest level of the subject and makes a to and fro motion. | |
| 4. Vocal | Demonstrator gives an audible call from its natural vocal repertoire. | Experimenter makes a gesture with slightly folded right palm, opening and closing the fingers gently at the chest level of the subject. | Any vocalisation from the repertoire of the subject is accepted. |
| 5. Flap Wings | Demonstrator flaps both wings with force while sitting on the perch. | Experimenter waves with her right palm in a to and from motion in front of the subject. | Flapping of both wings while sitting upright on a perch will be considered only. Flicking movement of wings usually associated with begging behaviour of juveniles will not be accepted. |

**Table S2**. Details of the subjects in the two experimental groups, target actions learnt including their learning order and the learning speed (total sessions taken and round).

| **Individual** | **Groups** | **Age (in years) / Sex (M/F)** | **Action demonstration order** | **Actions learnt (in order)** | **Total sessions taken (n$^{th}$ round)** |
|---|---|---|---|---|---|
| Marvel | **Test** | 2/F | Fluff | Lift leg | 3 (1$^{st}$) |
| | | | Spin | Vocal | 3 (1$^{st}$) |
| | | | Lift leg | Fluff | 7 (2$^{nd}$) |
| | | | Vocal | Spin | 14 (2$^{nd}$) |
| | | | Wings | Wings | 8 (2$^{nd}$) |
| Iron Man | **Test** | 1/M | Fluff | Fluff | 3 (1$^{st}$) |
| | | | Lift leg | Lift leg | 3 (1$^{st}$) |
| | | | Wings | Spin | 4 (1$^{st}$) |
| | | | Spin | Vocal | 3 (1$^{st}$) |
| | | | Vocal | Wings | 10 (2$^{nd}$) |
| Carrot | **Test** | 3/F | Lift leg | Lift leg | 3 (1$^{st}$) |
| | | | Fluff | Fluff | 2 (1$^{st}$) |
| | | | Wings | Vocal | 5 (1$^{st}$) |
| | | | Spin | Wings | 8 (2$^{nd}$) |
| | | | Vocal | Spin | 8 (2$^{nd}$) |
| Pickle | **Test** | 0.5/M | Spin | Spin | 2 (1$^{st}$) |
| | | | Lift leg | Lift leg | 2 (1$^{st}$) |
| | | | Vocal | Vocal | 9 (2$^{nd}$) |
| | | | Fluff | | |
| | | | Wings | | |
| Natasha | **Test** | 4/F | Lift leg | Lift leg | 6 (1$^{st}$) |
| | | | Spin | Spin | 4 (1$^{st}$) |
| | | | Vocal | Vocal | 9 (1$^{st}$) |
| | | | Wings | Fluff | 7 (1$^{st}$) |
| | | | Fluff | | |
| Thor | **Test** | 13/M | Fluff | Fluff | 3 (1$^{st}$) |
| | | | Lift leg | Spin | 6 (1$^{st}$) |
| | | | Wings | Lift leg | 12 (2$^{nd}$) |
| | | | Vocal | | |
| | | | Spin | | |

| | | | | | |
|---|---|---|---|---|---|
| Wanda (excluded) | **Test** | 15/F | NA | NA | NA |
| Morty | **Control** | 4/M | Spin | Lift leg | 8 (1st) |
| | | | Wings | Spin | 17 (2nd) |
| | | | Vocal | Fluff | 13 (2nd) |
| | | | Lift leg | | |
| | | | Fluff | | |
| Rick | **Control** | 4/M | Lift leg | Spin | 3 (1st) |
| | | | Spin | Fluff | 16 (2nd) |
| | | | Fluff | | |
| | | | Vocal | | |
| | | | Wings | | |
| Pepper | **Control** | 1/F | Lift leg | Spin | 10 (1st) |
| | | | Spin | Vocal | 2 (1st) |
| | | | Wings | | |
| | | | Fluff | | |
| | | | Vocal | | |
| Sherlock | **Control** | 13/M | Lift leg | Lift leg | 3 (1st) |
| | | | Fluff | Fluff | 11 (2nd) |
| | | | Spin | | |
| | | | Vocal | | |
| | | | Wings | | |
| Strange | **Control** | 1/M | Vocal | Spin | 19 (2nd) |
| **Was sent** | | | Fluff | Lift leg | 6 (2nd) |
| **to foster** | | | Spin | | |
| **home for 1** | | | Wings | | |
| **month after round 1** | | | Lift leg | | |

**Video S1.** Exemplary video of a *'spin'* test trial.

Three times the demonstrator responds to experimenter 1's hand command for *'spin'* by performing a *'spin'* followed by an immediate clicker and a reward. Following the three reinforced demonstrations, the test group subject on the right side receives the same hand command by experimenter 2, which is continued until the subject responds correctly and receives a clicker and subsequent reward too.

**Video S2.** Exemplary video of a *‘lift leg’* test trial.

Three times the demonstrator responds to experimenter 1’s hand command for *‘lift leg’* by performing a *‘lift leg’* followed by an immediate clicker and a reward. Following the three reinforced demonstrations, the test group subject on the right side receives the same hand command by experimenter 2, which is continued until the subject responds correctly and receives a clicker and subsequent reward too.

**Video S3.** Exemplary video of a *‘fluff’* test trial.

Three times the demonstrator responds to experimenter 1’s hand command for *‘fluff’* by performing a ‘fluff’ followed by an immediate clicker and a reward. Following the three reinforced demonstrations, the test group subject on the right side receives the same hand command by experimenter 2, which is continued until the subject responds correctly and receives a clicker and subsequent reward too.

**Video S4.** Exemplary video of a *‘vocal’* test trial.

Three times the demonstrator responds to experimenter 1’s hand command for *‘vocal’* by performing a *‘vocal’* followed by an immediate clicker and a reward. Following the three reinforced demonstrations, the test group subject on the right side receives the same hand command by experimenter 2, which is continued until the subject responds correctly and receives a clicker and subsequent reward too.

**Video S5.** Exemplary video of a *‘flap wings’* test trial.

Three times the demonstrator responds to experimenter 1’s hand command for *‘flap wings’* by performing a *‘flap wings’* followed by an immediate clicker and a reward. Following the three reinforced demonstrations, the test group subject on the right side receives the same hand command by experimenter 2, which is continued until the subject responds correctly and receives a clicker and subsequent reward too. The subject shows spontaneous imitation after it had been rewarded for the correct response.

**Video S6.** Exemplary video of a control trial *('vocal')*.

No conspecific demonstrator is present in the adjacent experimental room. The control group subject receives the hand command for '*vocal*', which is continuously repeated until the bird either shows the target action and is rewarded or 12sec have passed (shown here).

**Video S7**. Exemplary video of a control trial *('lift leg')* with social facilitator present.

In round 2, experimenter 1 stands in front of a 'social facilitator' bird in the adjacent experimental room to the left, to control for social enhancement. The control group subject on the right receives the hand command for '*lift leg'*, which is continuously given until the bird either shows the target action and is rewarded or 12sec have passed (shown here).
